# Discrepancies between observed data and predictions from mathematical modelling of the impact of screening interventions on *Chlamydia trachomatis* prevalence

**DOI:** 10.1101/389387

**Authors:** Joost Smid, Christian L. Althaus, Nicola Low

## Abstract

Mathematical modelling studies of *C. trachomatis* transmission predict that interventions to screen and treat chlamydia infection will reduce prevalence to a greater degree than that observed in empirical population-based studies. We investigated two factors that might explain this discrepancy: partial immunity after natural infection clearance and differential screening coverage according to infection risk. We used four variants of a compartmental model for heterosexual *C. trachomatis* transmission, parameterized using data from England about sexual behaviour and *C. trachomatis* testing, diagnosis and prevalence, and Markov Chain Monte Carlo methods for statistical inference. A model in which partial immunity follows natural infection clearance and the proportion of tests done in chlamydia-infected people decreases over time fitted the data best. The model predicts that partial immunity reduced susceptibility to reinfection by 72% (95% Bayesian credible interval 57-86%). The estimated screening rate was 4.6 (2.6-6.5) times higher for infected than for uninfected women in 2000; this decreased to 2.1 (1.4-2.9) in 2011. Other factors not included in the model could have further reduced the expected impact of screening. Future mathematical modelling studies investigating the effects of screening interventions on *C. trachomatis* transmission should incorporate host immunity and changes over time in the targeting of screening.

## INTRODUCTION

There is ongoing debate about the evidence to support screening for *Chlamydia trachomatis* (chlamydia) infection to reduce prevalence.^1-3^ *C. trachomatis* is the most commonly reported bacterial sexually transmitted infection (STI) in high-income countries; in 2016, about 128,000 cases of *C. trachomatis* were diagnosed among young people aged 15-24 years in England^4^ and over 1 million in the United States of America ^5^. *C. trachomatis* can cause pelvic inflammatory disease (PID) in women, which can lead to ectopic pregnancy and tubal factor infertility.^6^ However, *C. trachomatis* infection is often asymptomatic, so screening has been promoted to detect and treat asymptomatic infection to prevent reproductive tract morbidity and reduce transmission. In England, screening for chlamydia increased considerably with the National Chlamydia Screening Programme (NCSP) in 2003. Through the NCSP, free opportunistic screening is offered to sexually active women and men under 25 years of age, with nationwide roll-out achieved in 2008. Testing coverage in young women increased from 4% in 2000 to 35% in 2012.^7^ However, chlamydia prevalence, estimated in two cross-sectional population-based British National Surveys of Sexual Attitudes and Lifestyles (Natsal) was similar in adults aged 18 to 24 years; in 1999-2001 (Natsal-2), women 3.1% (95% confidence interval, CI 1.8-5.2) and men 2.9% (1.3-6.3) and in 2010-2012 (Natsal-3). Women 3.2% (2.2-4.6) and men 2.6% (1.7-4.0).^8^,^9^

Transmission dynamic modelling studies predict that screening at levels achieved by the NCSP in England should reduce *C. trachomatis* prevalence.^10^ These modelling studies describe sexual networks and the dynamics of infection transmission using different structures and levels of complexity.^11-14^ In a simpler model, without detailed sexual behaviour, Lewis and White inferred changes in *C. trachomatis* prevalence and incidence using time-series data about chlamydia testing and diagnoses in England between 2000-2015.^7,15^ Their model output proposed that prevalence had declined as chlamydia testing increased and increased as testing levels fell. An assumption in these models, irrespective of structure, is that amongst people without symptoms suggestive of infection, testing for chlamydia is not influenced by the underlying risk of infection in the screened population. In reality, the NCSP in England, and other chlamydia screening interventions, primarily test people at an increased risk of chlamydia.^16^ Further, with increasing chlamydia test coverage and falling test positivity^7^, if prevalence stayed at similar levels then the proportion of tests done in those at lower risk of infection must have increased.

Immunity also affects the model-predicted impact of screening if treatment inhibits the development of immunity otherwise experienced after natural clearance of infection.^17,18^ In a model accounting for immunity, individuals that clear infection naturally are temporarily or partially protected from the force of infection which results in a less rapid turnover of *C. trachomatis* within the modelled population. This reduces the predicted impact of screening. Immunity is often not included as part of the natural history in *C. trachomatis* transmission models^10^ and, in practice, not much is known about the strength and duration of immunity. Clinical and animal studies suggest that immunity is probably partial instead of fully protective.^19,20^ In one modelling study, Johnson and colleagues estimated a period of immunity of 6-17 years by fitting their model to chlamydia notification data, but they assumed that immunity was fully protective.^18^

In this paper, we use data about sexual behaviour and the prevalence of *C. trachomatis* in the general population of Great Britain from Natsal-2 and Natsal-3 and time-series data about chlamydia testing and diagnoses in England and across the same time period. Using a *C. trachomatis* transmission model, we investigated two hypotheses about factors that might attenuate the effects of a chlamydia screening intervention: the existence of long-lasting partial immunity; and differential chlamydia test coverage according to the risk of being infected.

## METHODS

### Data

Natsal is conducted by face-to-face and computer-assisted questionnaire amongst a stratified random sample of the resident population of Great Britain at ten year intervals since 1990.^21^ Natsal-2 includes data about 12,110 respondents aged 16-44 years from 1999-2001, and Natsal-3 includes data about 15,162 respondents aged 16-74 years from 2010-2012. ^22,23^ Starting with Natsal-2, a random sample of participants who have ever had sexual intercourse has been invited to provide a first catch urine sample, which is tested for the presence *of C. trachomatis* using a nucleic acid amplification test (3,608 respondents aged 18-44 years in Natsal-2 and a 4,550 respondents aged 16-44 years in Natsal-3). We used data from heterosexual respondents between 16-44 years about the number of new heterosexual partners in the last year, the respondent’s age at first heterosexual intercourse and the respondent’s age and partner ages at the time of first sexual intercourse with the first, second and third most recent heterosexual partner. We aggregated these data for both surveys because there were no significant differences for these variables between the datasets.^22^

We used estimates for the numbers of chlamydia tests and diagnoses from 2000 to 2011 in England from Chandra and colleagues.^7^ In this study, available data from several monitoring and surveillance systems in England, including NCSP, were collated to construct plausible minimum and maximum estimates for the numbers of tests and diagnoses each year for men and women in five-year age groups: 15-19, 20-24, 25-34 and 35-44 year olds. Chlamydia testing data did not distinguish between tests provided to people with symptoms suggestive of infection with *C. trachomatis* and screening tests amongst people without symptoms.

### Chlamydia transmission model

We developed a mathematical model to describe heterosexual *C. trachomatis* transmission in England from 2000 to 2011. The model uses differential equations and is described in detail in the Supplementary Information, part I; a brief summary is provided here. Model compartments and transmission rates are shown in Fig. 1 and Table 1.

**Figure 1:**
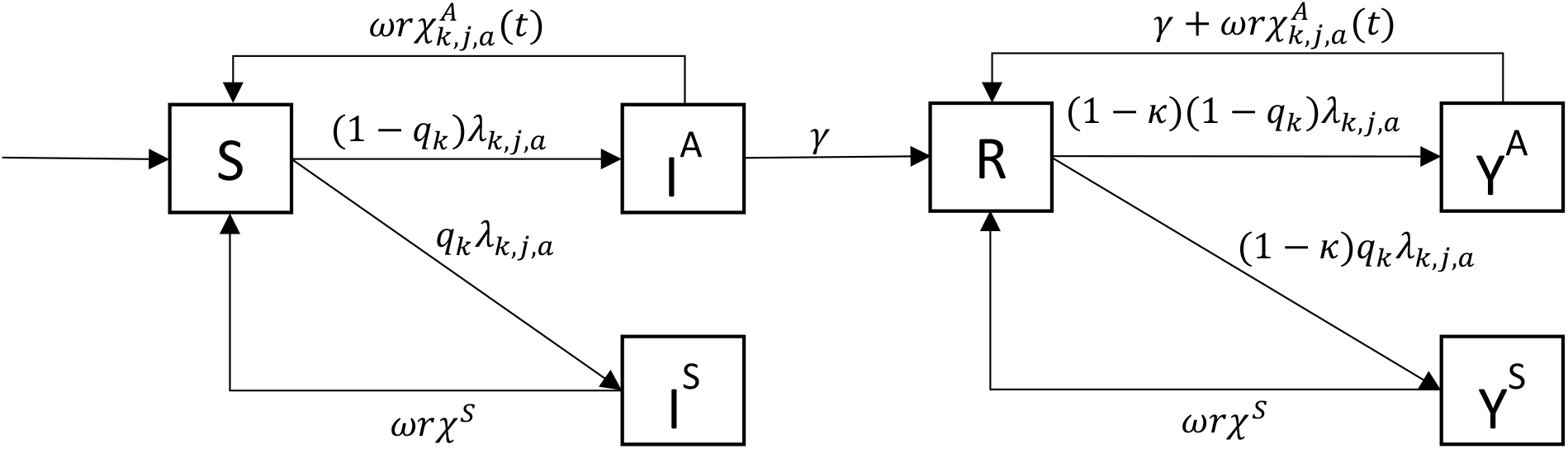
*Schematic illustration of the C. trachomatis transmission model. S susceptible; I^A^ asymptomatically infected; I^S^ symptomatically infected; R partially immune; Y^A^ asymptomatic reinfection; Y^S^ symptomatic reinfection. Parameters are described in Table 1*.

**Table 1:** *Description of parameters used in the transmission model and their values or prior distributions*

| Parameter | Description | Value/<br>Prior | Source |
| --- | --- | --- | --- |
| **Fixed parameters** |  |  |  |
| $c_{k,j,a}$ | Heterosexual partner change rate per year | See SI | 21,24 |
| $\rho_{k,a,a'}$ | Partner age distribution | See SI | 21,24 |
| $p_{k,a}$ | Probability of having had sex for the first time in the previous year | See SI | 21,24 |
| $f_{k,j}$ | Proportion of people in high activity class | See SI | 21,24 |
| $\alpha$ | Aging rate per year | 1 | - |
| $m$ | Redistribution rate between activity classes | 1 | 25 |
| $1/\gamma$ | Duration of the asymptomatic period without screening (years) | 433/365 | 17 |
| $\omega$ | Probability of successful treatment | 0.921 | 39 |
| $r$ | Probability of not being reinfected by a partner shortly after treatment | 0.806 | 40 |
| $1/\chi^S$ | Duration of the symptomatic period (years) | 33/365 | 41 |
| $\tau_{k,a,y}$ | Number of tests offered to people in class $(k, a)$ | See SI | 7 |
| **MCMC inferred parameters** |  |  |  |
| $\beta$ | Per partnership transmission probability | Beta(1,1) | |
| $\epsilon$ | Sexual mixing coefficient | Beta(1,1) | |
| $q_M$ | Fraction of symptomatic infections in men | Beta(1,1) | |
| $q_F$ | Fraction of symptomatic infections in women | Beta(1,1) | |
| $\eta_1$ | Parameter 1 for per capita ratio of screening between infected and susceptibles | U[1,10] | |
| $\eta_2$ | Parameter 2 for per capita ratio of screening between infected and susceptibles | Exp(U[-2,1]) | |
| $\kappa$ | Factor by which susceptibility to subsequent infection is reduced through partial immunity | Beta(1,1) | |

The model describes the transmission of chlamydia among susceptible people (*S*), infected people with a primary infection (*I*), people that have recovered naturally (*R*) and people with a repeated infection (*Y*). People are further stratified by sex *k* (men, women), age *a* (15-17, 18-19, 20-24, 25-34 and 35-44 years old) and sexual activity class *j* (low, high, defined by the average number of new heterosexual partners per year). Detailed age mixing behaviour and mixing between activity classes is modelled using methods proposed in a previous publication, where we accounted for differences in sexual behaviour reported by men and women.^24^ People can switch between activity classes at a rate that is proportional to the size of these classes.^25^ There is a time-dependent force of infection *λ_k,j,a_* that involves the per partnership transmission probability (*β*) and the heterosexual mixing patterns between ages and sexual activity classes. A fraction *q* of all new infections is symptomatic, the remainder being asymptomatic. People in the recovered compartment are partially immune with a reduced susceptibility to reinfection.

We assumed that all symptomatically infected people receive a chlamydia test at rate *χ*^5^ and are subsequently treated. After treatment, symptomatically infected people become susceptible again at rate *ω r χ^S^*, where 1 – *ω* is the probability of treatment failure, 1 – *r* the probability of being reinfected by a partner immediately after successful treatment and 1/*χ^S^* the average duration until treatment. Asymptomatically infected people can clear their infection naturally at rate *γ*, or are screened at a sex- and age-dependent rate 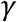 that depends on time t. The chlamydia screening rate amongst asymptomatic individuals differs according to the risk of being infected, which we refer to as differential screening coverage. In the model, we define differential screening coverage as the ratio of the screening rates in asymptomatic individuals who are infected with *C. trachomatis* and asymptomatic individuals who are uninfected. Arguably, this ratio decreases until 2011, as more screening is offered and becomes less targeted. We model differential screening coverage by a time-dependent parameter *η_k,a_*(*t*) that decreases exponentially as a function of the total yearly number of screening tests *Ξ_k,a_*(*t*):

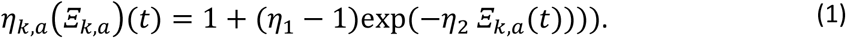

*η_k,a_*(*Ξ_k,a_*) starts at *η*_1_ and converges to one as the number of tests *Ξ_k,a_* becomes large, reflecting the situation in which screening is distributed homogeneously among all people. We then calculated the screening rate in asymptomatically infected people, 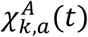, from *Ξ_k,a_*(*t*) and *η_k,a_*(*t*) by dividing the total number of screening tests in a given year by the total number of people eligible to receive screening, accounting for the differential screening coverage between asymptomatically infected and uninfected people:

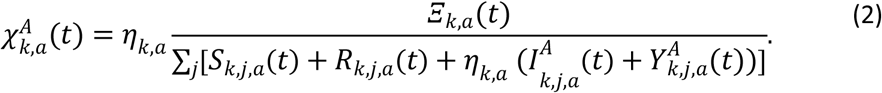

### Parameter inference

We inferred the values of certain parameters using Markov Chain Monte Carlo (MCMC) sampling (Table 1).^26^ We ran five separate MCMC chains for 10,000 MCMC steps. We assumed a binomial likelihood for the prevalence data and a negative binomial (NB) likelihood for the diagnoses data (Supplementary Information, part II). We checked convergence of the MCMC chains by computing the Gelman-Rubin convergence diagnostic.^27^ We used fixed values for the sexual behaviour parameters and some of the other parameters (*γ*, *r* and *χ^S^*) (Table 1). To explore the sensitivity of the model outcomes to parameters *γ*, *r* and *χ^S^*, we varied their values by 10% and ran the same MCMC algorithm again.

### Comparison of model variants

We used the chlamydia transmission model to investigate whether long-lasting partial immunity and differential screening coverage can explain discrepancies between model-predicted effects of screening and observed data in England. We considered four different model variants (Table 2). In models 1 and 2 we assumed no partial immunity (by fixing *κ* to zero), whereas models 3 and 4 can account for immunity (0 < *κ* ≤ 1, estimated using MCMC). In models 1 and 3, we assumed a differential screening coverage that does not change as screening increases in time, i.e. (*η*_1_ estimated using MCMC and fixing *η*_2_ to zero). In models 2 and 4 we allowed for a differential screening coverage that changes as screening increases (*η*_1_, e and *η*_2_ both estimated using MCMC). We compared the goodness of fit of the different model variants using the deviance information criterion (DIC). ^28,29^

**Table 2:** *Summary of parameters (mean and 95% CI of posterior distributions) for different models. The last two rows show the fit statistics of the models. *Kept as fixed values in these models*.

| Parameter | Model 1 | Model 2 | Model 3 | Model 4 |
| --- | --- | --- | --- | --- |
| $\beta$ | 0.61(0.57,0.65) | 0.6(0.56,0.63) | 0.85(0.79,0.92) | 0.84(0.77,0.91) |
| $\epsilon$ | 0.75(0.55,0.94) | 0.81(0.62,1) | 0.79(0.61,0.97) | 0.84(0.67,1.01) |
| $q_M$ | 0.13(0.09,0.18) | 0.09(0.02,0.16) | 0.14(0.09,0.18) | 0.1(0.04,0.16) |
| $q_F$ | 0.14(0.07,0.21) | 0.1(0,0.19) | 0.14(0.07,0.21) | 0.1(0.02,0.19) |
| $\eta_1$ | 2.06(1.36,2.76) | 6.05(2.28,9.83) | 2.51(1.68,3.34) | 6.04(2.69,9.38) |
| $\eta_2$ | 0* | 1.20 (0.19,7.59) | 0* | 0.98(0.42,2.29) |
| $\kappa$ | 0* | 0* | 0.7(0.57,0.84) | 0.72(0.57,0.86) |
| Log likelihood | -781(-783,-778) | -776(-780,-773) | -764(-767,-761) | -760(-763,-757) |
| DIC | 1431 | 1424 | 1401 | 1389 |

Model code (available on request) was implemented using R version 3.4.0.

## RESULTS

The posterior probability distributions for the model-estimated parameters were different for each model variant and resulted in a different goodness of fit (DIC) to the empirical data on *C. trachomatis* prevalence and diagnoses (Table 2). Comparing the DIC values allowed us to investigate the validity of the assumptions distinguishing the four model variants. A model including a decreasing per capita ratio of screening tests in infected compared to susceptible people and partial immunity (model 4) described the data better (lowest DIC) than models where the composition of the screened population was assumed fixed in time or without partial immunity (Table 2). A model including partial immunity but no changes over time in the composition of the screened population (model 3) fitted second best to the data with a 12 points’ higher DIC value. Models without partial immunity fitted considerably worse to the data. The model fit and the posterior distributions showed limited sensitivity to the exact values of the fixed parameters *γ*, *r* and *χ^S^* (Supplementary Information, part III). In all models, the MCMC chains converged well (Gelman-Rubin diagnostic below 1.1).

Partial immunity decreased susceptibility to reinfection after natural clearance of a first infection by 72% (95% Bayesian credible interval (CrI) 57%-86%) in model 4 (Table 2). The estimate of partial immunity in the second-best fitting model 3 (fixed per capita ratio of tests in infected compared to susceptible people) was similar (70%, 95% CrI 57%-84%).

Fig. 2 shows the posterior prevalence predicted by the best-fitting model 4 and the observed levels of *C. trachomatis* prevalence in England in 1999-2000 and 2010-2011. The uncertainty about the prevalence estimates from empirical data is large due to small sample sizes.^8,9^ The model predicted a drop in prevalence for women between 15-24 years of 51% (95% CrI 47%-54%) and for men between 15-24 years of 56% (95% CrI 51%-60%), also see Fig. 5. The mean model-predicted prevalence falls within the confidence intervals of the prevalence data for all age groups and both periods, with the exception of women aged 18-19 years in 2011. However, some inconsistencies remain between the observed and model-predicted chlamydia prevalence. In particular, for women the mean model-predicted prevalence is higher than the mean prevalence from data for 2000 and lower for 2011 for almost all age groups.

**Figure 2:**
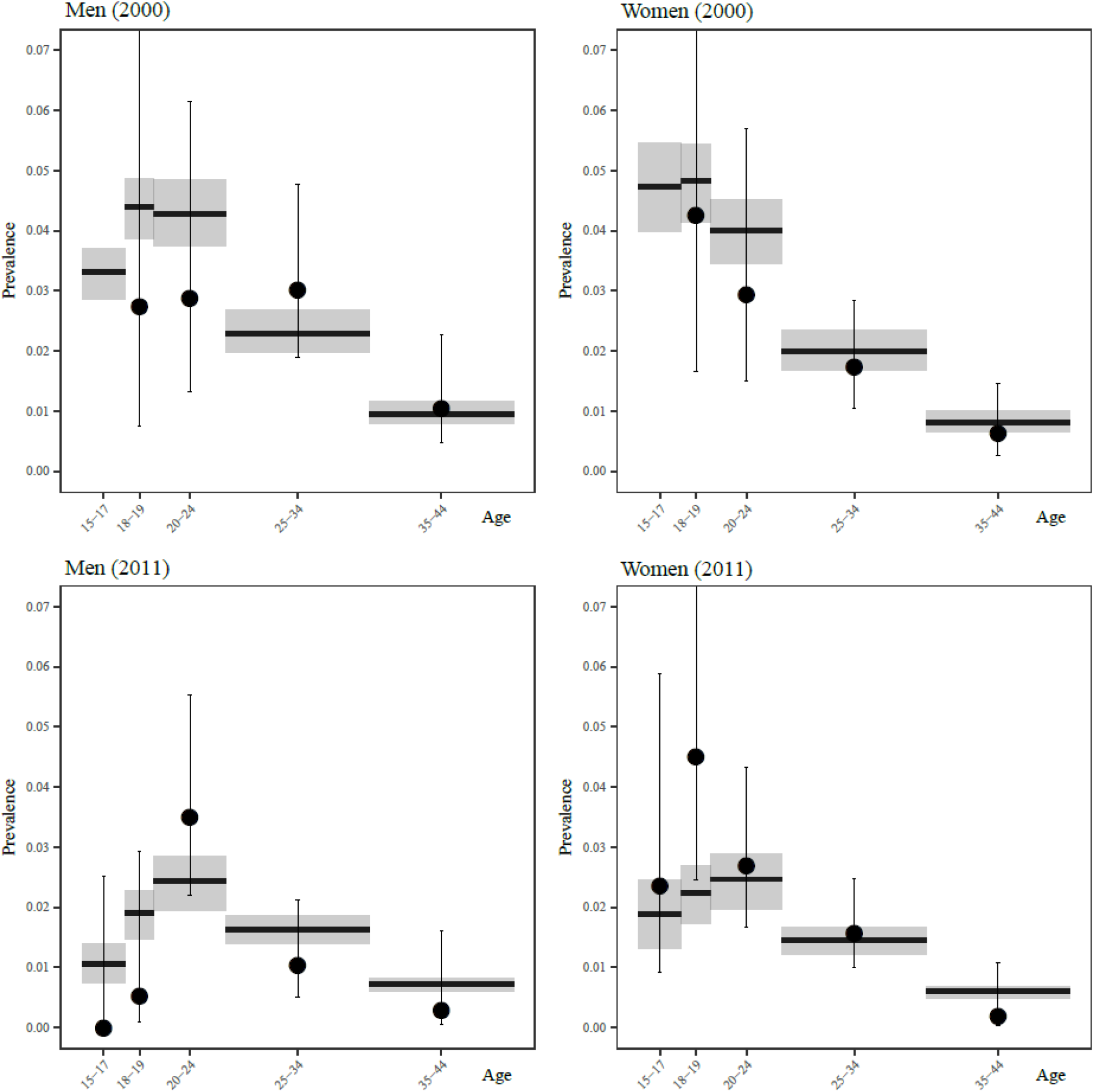
*Model fit to age-specific chlamydia prevalence for men and women in 2000 and 2011. Black dots and vertical bars: Estimated prevalence from Natsal (mean and 95% confidence intervals). Grey boxes: Full model (model 4) including changes in the proportion of tests done in infected individuals and partial immunity (posterior mean and 95% Bayesian credible intervals).*

Empirical data show that the per capita number of diagnoses of *C. trachomatis* infections has increased between 2000 and 2011 (Fig. 3). Although model 4 could roughly reproduce this trend, some differences remain between the model fit and the data. In general, the model shows a less steep increase in the number of diagnoses from 2000 to 2011 for men and women between 20-24 years than was estimated by Chandra and colleagues.^7^ For men aged 15-19 years, the model-predicted number of diagnoses increased more steeply than observed from data.

**Figure 3:**
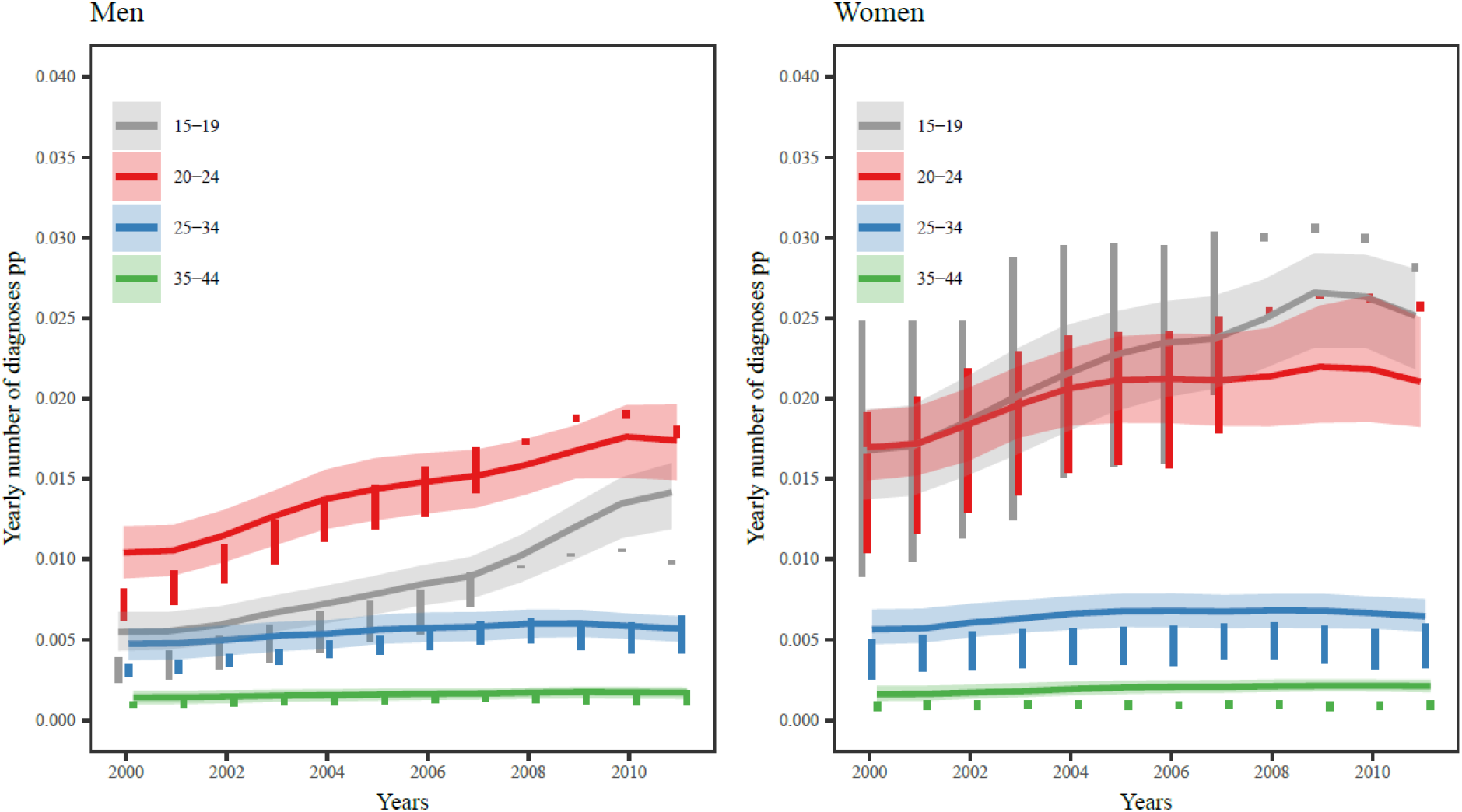
*Model fit to age-specific per capita number of diagnoses for men and women between 2000 and 2011. Vertical bars: Minimum and maximum estimates for number of diagnoses from Chandra and colleagues.^7^ Coloured lines and shaded areas: Full model (model 4) including changes in the proportion of tests done in infected individuals and partial immunity (posterior mean and 95% Bayesian credible intervals*).

In model 4, the expected screening rate was 4.6 (95% CrI 2.6-6.5) times higher for infected than for uninfected women in 2000; this decreased to 2.1 (95% CrI 1.4-2.9) in 2011. For men, this ratio decreased from 5.5 (95% CrI 2.7-7.9) in 2000 to 3.2 (95% CrI 2.0-4.4) in 2011 (Fig. 4). The Bayesian credible intervals around these estimates are large. For men and women aged 25-44 years, we found no decrease in the ratio. The decrease in differential screening coverage in the model is the consequence of increased testing volume and not of decreased chlamydia prevalence, but it does affect prevalence. To show this, we investigated what would have happened to the predicted chlamydia prevalence if the estimated differential screening coverage as estimated for 2000 is maintained, while increasing the total number of screening tests as in the data. We used the estimated posterior distribution for *η*_1_ from model 4, while fixing *η*_2_ to zero. In this hypothetical scenario, the predicted decrease of chlamydia prevalence was significantly higher than the decrease predicted by model 4 (Fig. 5).

**Figure 4:**
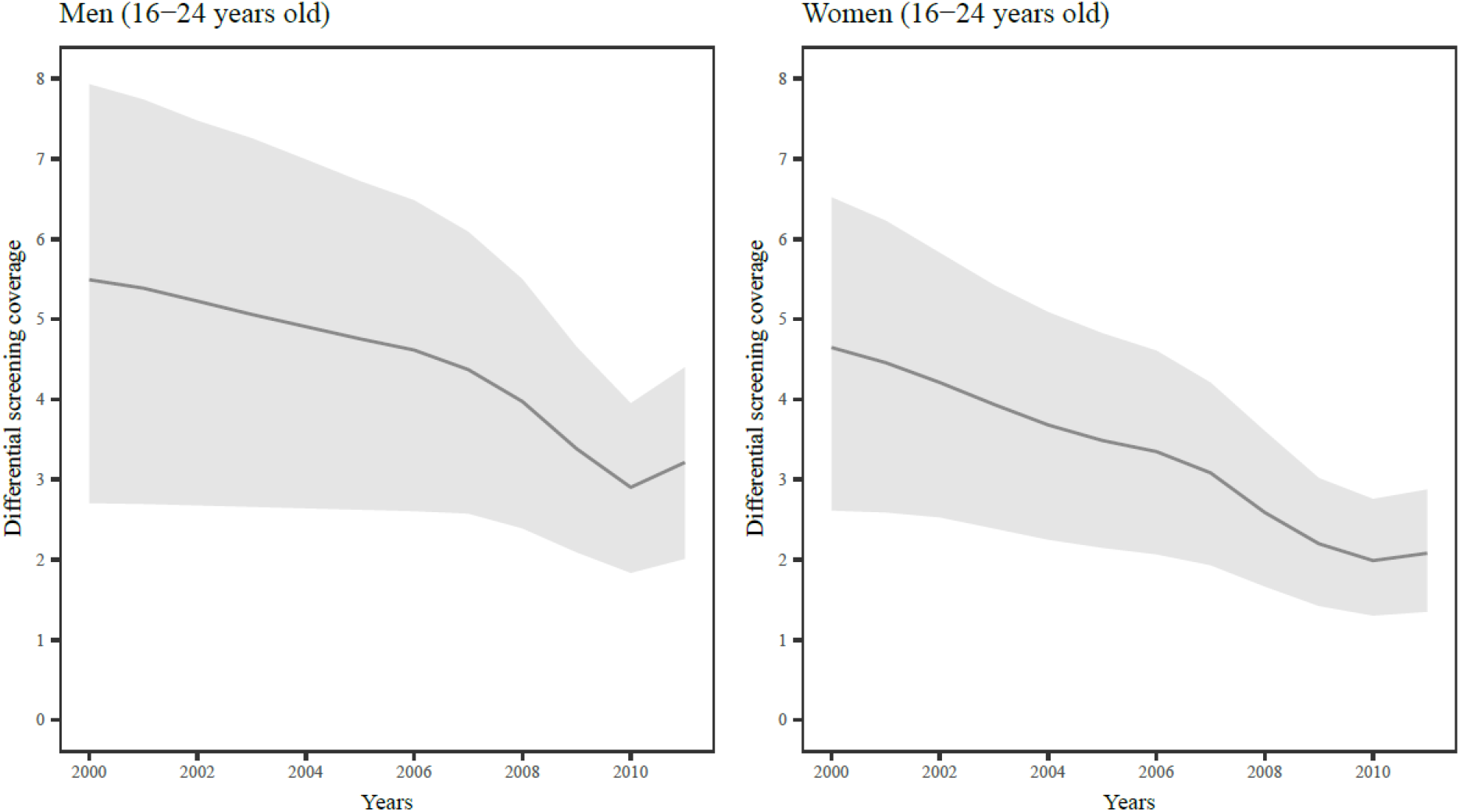
*Differential screening coverage: Model-estimated change in the ratio of the screening rates in infected vs. susceptible individuals for men and women of 16-24 years old between 2000 and 2011. Coloured lines and shaded areas: Posterior mean and 95% Bayesian credible intervals*.

**Figure 5:**
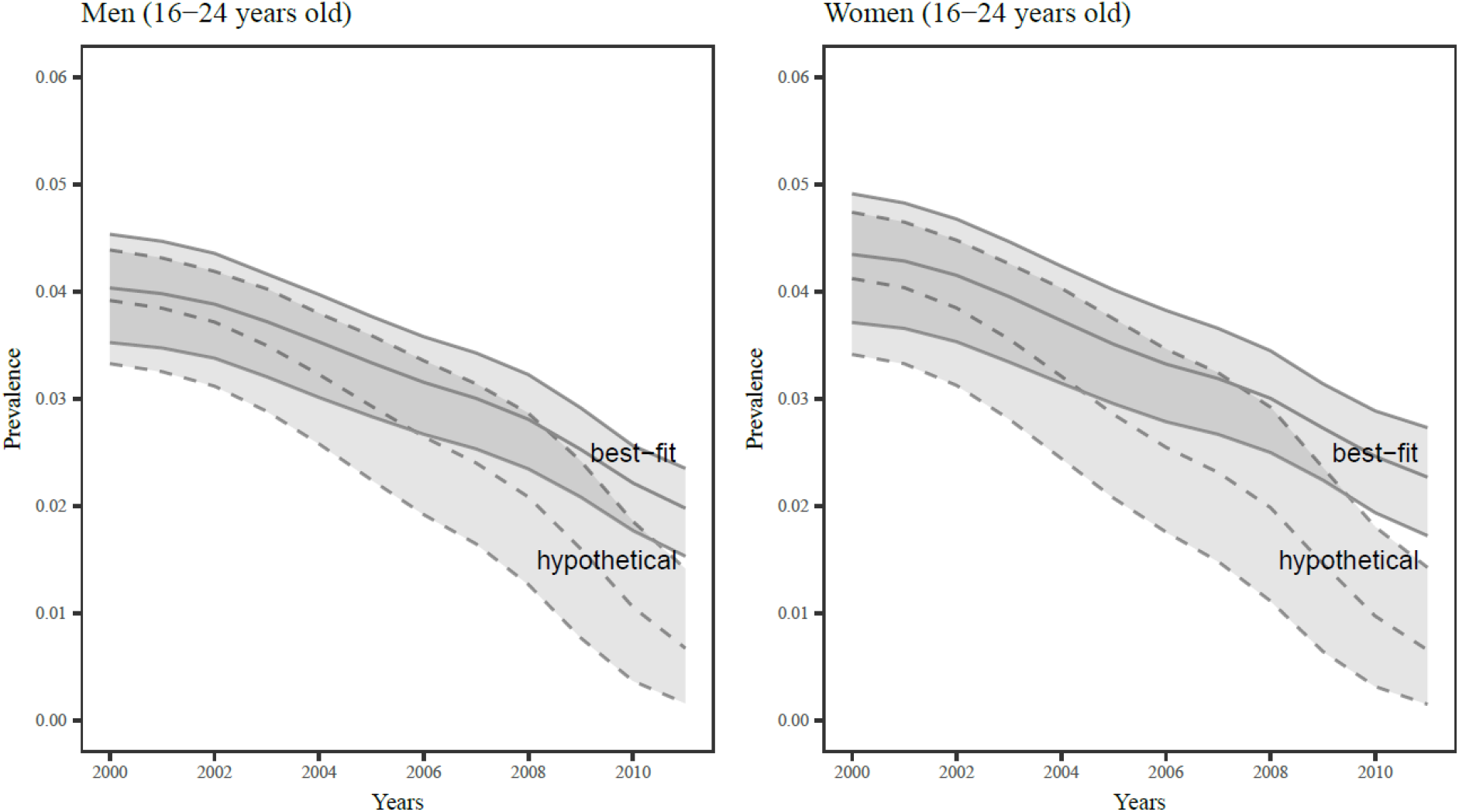
*Model-estimated change in chlamydia prevalence in men and women between 2000 and 2011. Best-fit: Full model (model 4) including changes in the proportion of tests done in infected individuals and partial immunity (posterior mean and 95% Bayesian credible intervals). Hypothetical: Scenario in which the proportion of screening tests in infected individuals is kept at the same level as was estimated for 2000*.

## Discussion

In this study, we compared four model variants in a compartmental model of *C. trachomatis* infection dynamics that includes partial immunity after natural clearance of infection and changes in differential screening coverage over time. The model variant that fitted best to empirical data about *C. trachomatis* prevalence and diagnoses in England between 2000 and 2011 (model 4) included both partial immunity (a reduction of susceptibility of 56%-82%) and a decreasing proportion of screening tests done in infected people over time. Although both factors attenuate the model-predicted effectiveness of screening, the model including both factors still predicted a decrease of chlamydia prevalence between 2000 and 2011 of 50-60% in the age groups targeted for screening.

A strength of this study is the availability of time-series data about chlamydia testing and diagnosis data,^7^ and repeated cross-sectional surveys of *C. trachomatis* prevalence and sexual behavioural data from the same population over the same time period.^22,23^ The testing data avoided the need, in other modelling studies, for strong assumptions about (unobserved) levels of screening coverage.^11,13,30^ Second, because of the importance of age as a risk factor for chlamydia, we included detailed age structure and age-dependent sexual behaviour in our model, which allowed us to estimate chlamydia prevalence in different age groups and fitted these to data about age-specific prevalence and diagnoses.^24^ We made optimal use of age-specific data to estimate the values about the unknown model parameters, including those quantifying immunity and the distribution of screening tests. Third, by synthesizing the data in a Bayesian framework we could quantify the distinct effects of partial immunity and dynamic changes in the distribution of screening tests, whilst accounting for uncertainty about the model parameters.

Our study has also limitations. First, the testing and diagnosis data, rather than the chlamydia prevalence data mainly drove the Bayesian inference in our model. Prevalence data added less to the total likelihood because the surveys included a small sample of the target population for two periods only (1999-2001 and 2010-2012), whereas the data about diagnoses are from the whole of England for each year between 2000-2011. To down weight the diagnosis data compared with the prevalence data, we modelled their likelihood using a negative binomial distribution instead of a Poisson distribution, which introduced a moderate dispersion in the diagnoses data, reflecting additional bias that we could not model explicitly. Second, the assumption of an exponential relationship between the total number of tests *T* in a given year and the per capita ratio of testing in infected compared with susceptible people *η*(*T*) in that year might be an over-simplification. An exponential function makes sense because convergence to one as the number of tests becomes large indicates the increasingly equal distribution of tests among all sexually active people but another, non-continuous function might be more suitable, particularly when screening policy has changed over time.^31^ Finally, we assumed only one level of partial immunity, which is a likely over-simplification, given the possibility of heterogeneity in immune response at the level of the individual or strain-specific immunity, and potential effects of repeat infection. It is unclear how these assumptions and levels of heterogeneity would affect our results.

The results of our study suggest that mathematical models that neglect changes over time in the distribution of screening tests among modelled subpopulations or do not include protective immunity after natural clearance of *C. trachomatis^10-14^* over-estimate the effects of screening interventions on prevalence. However, our model did not fully reconcile the model predictions with the data either, with a larger model-predicted decrease of *C. trachomatis* prevalence between 2000-2011 than expected from data.^9^ According to our model, screening and treating *C. trachomatis-infected* people resulted in a reduction in prevalence even when accounting for immunity. This finding does not support the results of a modelling study that suggested that widespread testing and treatment diminish population level immunity and can result in an increase in incidence and prevalence.^32^ Other factors have been proposed as reasons for the similarity of estimated *C. trachomatis* prevalence in the Natsal-2 and Natsal-3 surveys. First, increases in sexual risk-taking behaviour could have resulted in increased transmission of *C. trachomatis*, countering the effects of screening. However, amongst the sexual behaviours measured in Natsal-2 and Natsal-3, the only difference among heterosexual adults was a slight decrease in the proportion of men with multiple partners with whom no condoms was used.^22^ We do not believe that this change would be sufficient to abolish the model-predicted reduction in prevalence and we did not investigate this possibility because our model did not explicitly include condom use. Second, a temporary reduction in *C. trachomatis* prevalence could have occurred during the period between the Natsal surveys. In the modelling study by Lewis and White, which also used the data on chlamydia testing and diagnoses reported by Chandra and colleagues, model-inferred prevalence decreased until 2008 and then increased back to baseline.^15^ That model did not take into account differential screening coverage over time and inferred large increases in incidence to balance the increase in screening rates.^33^ In theory, a reduction in the rate of successful partner notification could also limit the impact of a chlamydia screening intervention. However, in a study that compared a pair model including partner notification with a simpler model without partnerships,^34^ changes in partner notification rates resulted in very modest changes in incidence. Third, it has been suggested that *C. trachomatis* transmission is maintained by infection in the female rectum that is not adequately treated.^35,36^ We deemed an analysis of the role of anorectal infections beyond the scope of this study because of the uncertainty about autoinoculation probabilities.

Our findings have implications for future research and practice. First, discrepancies remain between the observed estimates of *C. trachomatis* prevalence and the model predictions, even in our best-fitting model. Future modelling studies should investigate other factors, such as those listed above, which could bridge the remaining gap between data and the model predictions. Second, clinical immunological studies that can provide more information about the strength and duration of immunity following natural clearance of *C. trachomatis* would be very valuable. Third, our model predicted a larger reduction in *C. trachomatis* prevalence if the ratio of screening coverage in infected compared with uninfected individuals remained high. This finding has implications for defining targets for screening performance. After 2011 in the NCSP in England, targets for screening coverage were replaced by indicators for diagnosis rates to maintain high levels of diagnosed infections for a given test volume.^37^ The optimal diagnosis rate and its relationship with population prevalence, however, remains unknown.^38^

In conclusion, partial immunity against reinfection and changes in differential screening coverage over time might have limited the reduction in *C. trachomatis* prevalence that would be expected for the level of screening coverage achieved in England. The full range of factors that account for discrepancies between the observed prevalence data and model-predictions has not been elucidated. Future mathematical modelling studies that aim to investigate the effects of screening interventions on *C. trachomatis* transmission should incorporate host immunity and changes over time in the targeting of screening.

## ACKNOWLEDGEMENTS

JHS received support from the Swiss National Science Foundation (grant number 160320). CLA received support from the LUSTRUM Program of Research, funded by the National Institute for Health Research (NIHR) under its Program Grants for Applied Research Program (Reference Number RP-PG-0614-20009).

## AUTHOR CONTRIBUTIONS

JHS implemented and analysed the model. JHS wrote the initial draft of the manuscript. CLA and NL reviewed and commented on the manuscript.

